# Electrophysiological indices of anterior cingulate cortex function reveal changing levels of cognitive effort and reward valuation that sustain task performance

**DOI:** 10.1101/199687

**Authors:** Akina Umemoto, Michael Inzlicht, Clay B. Holroyd

## Abstract

Successful execution of goal-directed behaviors often requires the deployment of cognitive control, which is thought to require cognitive effort. Recent theories have proposed that anterior cingulate cortex (ACC) regulates control levels by weighing the reward-related benefits of control against its effort-related costs. However, given that the sensations of cognitive effort and reward valuation are available only to introspection, this hypothesis is difficult to investigate empirically. We have proposed that two electrophysiological indices of ACC function, frontal midline theta and the reward positivity (RewP), provide objective measures of these functions. To investigate this issue, we recorded the electroencephalogram (EEG) from participants engaged in an extended, cognitively-demanding task. Participants performed a time estimation task for 2 hours in which they received reward and error feedback according to their task performance. We observed that the amplitude of the RewP, a feedback-locked component of the event related brain potential associated with reward processing, decreased with time-on-task. Conversely, frontal midline theta power, which consists of 4-8 Hz EEG oscillations associated with cognitive effort, increased with time-on-task. We also examined how these phenomena changed over time by conducting within-participant multi-level modeling analyses. Our results suggest that extended execution of a cognitively-demanding task is characterized by an early phase in which high control levels combine with strong reward valuation to foster rapid improvements in task performance, and a later phase in which high control levels counteract waning reward valuation to maintain stable task performance.

---

Goal-directed behavior often requires cognitive control in order to facilitate the execution of non-automatic behaviors (Norman and Shallice, 1986). It is believed that the application of control feels effortful (e.g., Botvinick and Braver, 2015; Shenhav et al., 2017) and that cognitive effort is inherently aversive, such that people tend to avoid cognitively demanding tasks (Kool et al., 2010; Inzlicht et al., 2015; McGuire and Botvinick, 2010; Westbrook and Braver, 2015). This process is thought to recruit a mechanism that weighs the benefits of applying control against the effort-related costs in doing so (Botvinick and Braver, 2015; Shenhav et al., 2017; Westbrook and Braver, 2015). For example, although prolonged cognitive effort normally results in mental fatigue, which disrupts task performance, these performance decrements can be counteracted if subjects are offered motivational incentives (Boksem et al., 2006; Hockey, 2011; Hopstaken et al., 2015; Lorist et al., 2005; Boksem and Tops, 2008, for review). Yet despite an upsurge of interest in this topic in recent years, the neurocognitive mechanisms that sustain cognitive effort are not well-understood.

## Anterior cingulate cortex function in reward processing and effortful control

Recently, several theories and computational models have proposed that anterior cingulate cortex (ACC) may provide such a mechanism (e.g., Botvinick and Braver, 2015, Holroyd and McClure, 2015; Holroyd and Umemoto, 2016; Holroyd and Yeung, 2012; Shenhav et al., 2013, 2017; Vassena et al., 2017; Verguts et al., 2015). The function of ACC is famously controversial (Ebitz and Hayden, 2016). Nevertheless, accumulating evidence suggests that the caudal subdivision of ACC, which is formally known as “anterior midcingulate cortex” (Vogt, 2009; see also Shackman et al., 2011), may serve as a computational hub that integrates cognitive processes related to motivation and control. For example, Holroyd and Yeung (2012) proposed that ACC is responsible for motivating and selecting extended action sequences based on learned task values. According to this view, instead of concerning itself with the minutia of moment-to-moment behaviors (such as typing each letter in a manuscript), ACC regulates behavior at a higher level of abstraction (such as whether or not to write the manuscript at all), and determines how well the task should be performed at a global level (cf. Botvinick et al., 2009; Botvinick, 2012). Holroyd and McClure (2015) later implemented these ideas in a computational model of rodent behavior. Although these theories differ in their specifics, they hold in common the idea that ACC regulates the control levels it invests in a task according to the rewards received for doing so.

A challenge in studying these neurocognitive processes, however, is that the sensations of reward valuation and effort expenditure are available only to introspection, rendering them difficult to assess empirically. Task performance is an imperfect proxy for these processes because participants can perform a task well either because it is easy or because they apply enough effort to make it appear to be easy. Therefore, objective measures of reward valuation and cognitive effort, were they available, would provide insight into how the control system self-regulates (Botvinick and Braver, 2015). Here we propose that electrophysiological correlates of cognitive effort and of reward valuation can fulfill this purpose, and investigate this proposal by recording the electroencephalogram (EEG) from subjects as they perform an extended, mentally-fatiguing task.

## Electrophysiological correlates of reward valuation and cognitive effort

We addressed this question by utilizing two electrophysiological correlates of ACC activity: the reward positivity (RewP) and frontal midline theta (FMT) oscillations (Holroyd and Umemoto, 2016). More commonly known as the feedback error-related negativity, the RewP is an event-related potential (ERP) component that is differentially modulated by feedback stimuli with negative vs. positive valence (Miltner et al., 1997); the component has recently been renamed the RewP because it appears to be relatively more sensitive to reward outcomes than to error outcomes (Holroyd and Umemoto, 2016). Holroyd and Coles (2002) proposed that the RewP is produced by fast, phasic midbrain dopamine reward prediction error signals modulating ACC activity. Consistent with the theory, numerous studies have confirmed that the RewP indexes a reward prediction error signal (Sambrook and Goslin, 2015; Walsh and Anderson, 2012; but see Janssen et al., 2016). Given the inverse problem, the neural source of the RewP is less clear, but the balance of evidence suggests that it is produced in anterior midcingulate cortex (e.g., Miltner et al., 1997; Becker et al., 2014; but see Proudfit, 2015). Of particular relevance to this study, RewP amplitude correlates positively with individual differences in reward sensitivity (Bress and Hajcak, 2013, Umemoto and Holroyd, 2017; see also Cooper et al., 2014; Liu et al., 2014; Parvaz et al., 2016), and negatively with individual differences in depression levels (e.g., Umemoto and Holroyd, 2017; Proudfit, 2015), which suggests that the RewP may index subjective levels of reward valuation (see Holroyd and Umemoto, 2016 for review).

FMT consists of 4 to 8 Hz EEG oscillations distributed over frontal-central regions of the human scalp. Source localization studies of FMT point to ACC as the neural generator (e.g., Asada et al., 1999; Cavanagh and Frank, 2014; Ishii et al., 1999; Luu and Tucker, 2001), and FMT power appears to reflect an effortful control process (e.g., Cavanagh and Frank, 2014; Holroyd and Umemoto, 2016). Notably, cognitive tasks elicit two kinds of related FMT signals: phasic and ongoing. Brief, phasic changes in FMT power are observed immediately following error commission, during periods of response conflict (e.g., Cavanagh and Frank, 2014; Luu and Tucker, 2001), and with subjective conflict and surprise in an economic decision-making task (Lin et al., 2017, August); one study reported that phasic FMT power decreased with time-on-task (Washer et al., 2014). By contrast, ongoing FMT power is observed over extended periods of task execution (e.g., Asada et al., 1999; Ishii et al., 1999), as well as during the off-task, resting state (Scheeringa et al., 2008). It is associated with sustained mental effort, as observed, for example, when participants perform arithmetic calculations (Hsieh and Ranganath, 2014; Mitchell et al., 2008). Furthermore, FMT power rises with time on task (e.g., Barwick et al., 2012; Paus et al., 1997; Wascher et al., 2014), suggesting that ACC contributes to sustaining effortful behavior in the face of growing mental fatigue.

## Present study

We have previously proposed that FMT reflects the control signal applied by ACC over task performance, and that RewP amplitude reflects the reward signal propagated to the ACC for the purpose of regulating the control level (Holroyd and Umemoto, 2016). In this way, ACC is well-positioned to motivate task performance (Holroyd and McClure, 2015; Holroyd and Yeung, 2012). Although previous studies have utilized these signals to study reward valuation and cognitive effort independently, to our knowledge none have examined how these processes together sustain behavior on a mentally fatiguing task.

Here we utilized RewP amplitude and FMT power to examine whether these signals reflect neural processes related to subjective reward valuation and effort expenditure, respectively, in subjects performing an extended, cognitively-demanding task. For this purpose, we recorded the EEG from participants while they performed a standard time estimation task for 2 hours (Miltner et al., 1997). In line with previous observations, we expected that FMT power would gradually increase with time-on-task (Barwick et al., 2012; Paus et al., 1997; Wascher et al., 2014). Further, we predicted that RewP amplitude would decrease as performance deteriorated due to growing mental fatigue. We also expected that these signals would provide independent sources of information on the role of ACC in sustaining task performance, although we did not have any specific predictions about how they would interact. Lastly, we explored whether ACC function expressed as a dimension of personality related to reward sensitivity and persistence, as we have recently proposed (Holroyd and Umemoto, 2016; see also Umemoto and Holroyd, 2016, 2017). For this purpose, participants answered several personality questionnaires related to motivation, reward sensitivity, and levels of depression, which we explored in relation to their behavior and to these electrophysiological signatures of ACC function.

## Materials and methods

### Participants

Sixty five undergraduate students were recruited from the University of Victoria. The sample size was determined based on past studies examining individual differences related to the RewP (e.g., Bress and Hajcak, 2013; Nelson et al., 2016). A sensitivity analysis using the G*power software program (Erdfelder et al., 1996) indicated sufficient statistical power to detect small effects for the main within-subject analyses^1^. Participants were recruited from the Department of Psychology subject pool to fulfill a course requirement or earn bonus credits. All subjects (16 males, 10 left-handed, age range=17-23 years, mean age = 19.3 +/- 1.5 years) had normal or corrected-to-normal vision. Each also received a monetary bonus in addition to the credits, the amount of which depended on their task performance (see below). All of the subjects provided informed consent as approved by the local research ethics committee. The experiment was conducted in accordance with the ethical standards prescribed in the 1964 Declaration of Helsinki.

### Task Design

We conducted a time estimation task that required participants to estimate 1 second on every trial (Miltner et al., 1997). Each trial started with a presentation of a visual cue in the form of a white central cross (1° by 1° square visual angle) on the computer screen. Participants were instructed to press a left button on a computer mouse using their right hand when they believed that 1 second had elapsed following the cue onset. After 600 ms following the response, they were presented with a feedback stimulus indicating whether their response was on-time or not on-time. The response was considered on-time if it occurred within a narrow window of time centered around 1 second, the size of which was adjusted from trial to trial according to a staircase procedure, as follows. At the start of the experiment the size of the window was initialized at 200 ms, such that responses occurring between 900 and 1100 ms were considered correct. The size of the time window (i.e., time-window size) was then adjusted depending on the participant’s performance: error responses caused the time window to increase by 10 ms (making the task easier), and correct responses caused the time-window to decrease by 10 ms (making the task harder). This manipulation ensured that participants received reward feedback on approximately 50% of the trials (Miltner et al, 1997). The feedback stimuli were represented by abstract symbols presented at fixation (3.3° by 3.3° square visual angle). For half of the participants, a white circle indicated that they earned 1 cent for that trial, and a white diamond indicated that they did not earn any reward for that trial. This mapping was reversed for the other half of the participants. After a variable inter-trial interval of 1400 ms, 1500 ms, or 1600 ms, determined at random, the next trial began with a presentation of the visual cue.

Further, we offered an additional motivational incentive to enhance individual differences in task performance by providing all of the participants an opportunity to participate in a lottery to win a CAN $100 Amazon.com gift card. Every participant earned at least 1 “ticket” regardless of their performance, which ensured that all subjects had at least a small chance of receiving the prize. In addition, the participants were told (truthfully) that the better performers would earn extra tickets. For this, when all of the participants’ data were collected, we calculated each participant’s mean time-window size across all trials and compared them against the grand mean time-window size averaged across all participants. Participants received an extra ticket for every 5 ms decrement with respect to the grand mean time-window size. For example, an individual average time-window size that was 15 ms less than the grand mean time-window size would earn that participant 4 tickets total (3 additional tickets plus the baseline 1 ticket). Upon completing the experiment, all of the tickets were entered into a metaphorical lottery box, from which we randomly selected 2 winning participants. The two winners were then contacted and received the award via email afterword.

### Task Procedure

Participants were seated comfortably in front of a computer monitor (1024 by 1280 pixels) at a distance of about 60 cm in a dimly lit room. The task was programmed in Matlab (MathWorks, Natick, MA, USA) using the Psychophysics Toolbox extension (Brainard, 1997; Pelli, 1997). Before the experiment was described to them, all of the participants read a form that explained the opportunity to win a CAN $100 Amazon gift card. Participants were told that if they decided to consent to participate in this opportunity, at least one ticket would be entered in their name into the “lottery”, and that additional tickets would be entered if their overall performance was better than the average performance across all of the participants. Consent was indicated by participants providing a contact email address on the form; only two participants did not provide consent.

Participants were then provided with both written and verbal instructions about the time estimation task. They practiced the task for 20 trials before starting the task proper, which consisted of 16 blocks of 95 trials each. Participants were not informed about the exact number of trials or blocks to complete, but instead were told that they would perform the task for about 2 hours. Self-paced between-block rest periods were provided, and after about 1 hour participants relaxed during a longer rest period while the experimenter checked the electrode impedances. Participants were told that the reward they accumulated across trials would be paid out to them at the end of the experiment, and that they should estimate 1 second on each trial as accurately as possible in order to maximize their reward earnings. Upon completing the experiment they answered several personality questionnaires (see below). These were followed by a post-experiment paper-and-pen questionnaire that asked about the participant’s overall experience of the experiment, the strategies they employed (if any), and their level of task engagement on a scale of 1 to 5, with 1 indicating not at all engaged and 5 indicating very engaged.

### Questionnaires

Participants completed a total of six personality questionnaires related to motivation, reward sensitivity, and depression symptoms, administered through LimeSurvey (https://www.limesurvey.org/) on the same computer where the task was performed. The personality questionnaires administered included 1) the Persistence Scale (Cloninger et al., 1993; Gusnard et al., 2003), which assesses the tendency to overcome daily challenges on a scale of 1 (definitely false) to 5 (definitely true). 2) The 8-item Reward Responsiveness (RR) Scale (Van den Berg et al., 2010), which is a self-report measure of reward-related behavior on a scale of 1 (strong disagreement) to 4 (strong agreement). (3) The 18-item Temporal Experience of Pleasure Scale (TEPS), which assess two components of hedonic capacity, namely consummatory pleasure (TEPS-C: i.e., “liking” or in-the-moment experience of pleasure) and anticipatory pleasure (TEPS-A: i.e., “wanting”), on a scale of 1 (“very false for me”) to 6 (“very true for me”) (Gard et al., 2006). 4) The 18-item Apathy Evaluation Scale (AES; Marin et al., 1991), which measures lack of motivation regarding the behavioral, cognitive, and emotional aspects of goal-directed behavior on a scale of 1 (very characteristic) to 4 (not at all characteristic). 5) The 22-item Ruminative Responses Scale (RRS: Treynor et al., 2003), which measures the propensity to ruminate in response to depressed mood on a scale of 1 (almost never) to 4 (almost always). 6) The 21-item short-form of the Depression Anxiety Stress Scale (DASS-21) (Lovibond and Lovibond, 1995), which measures severity of depression, anxiety, and stress on a scale from 0 (“did not apply to me at all”) to 3 (“applied to me very much, or most of the time”). However, the rumination scale and the anxiety and stress subscales of the DASS-21 were not included in the analyses as they tended to strongly correlate with other variables (for example, with the depression scores), and because they were not the primary focus of the study. Summed total scores were used for each of the questionnaires such that high scores indicated, respectively, high persistence, high reward responsiveness, high hedonic capacity (or reduced anhedonia), high levels of apathy, and high levels of depression.

### EEG Data Acquisition and Pre-processing

The EEG was recorded using a montage of 30 electrode sites in accordance to the extended international 10–20 system (Jasper, 1958). Signals were acquired using Ag/AgCl ring electrodes mounted in a nylon electrode cap with an abrasive, conductive gel (EASYCAP GmbH, Herrsching-Breitbrunn, Germany). Signals were amplified by low-noise electrode differential amplifiers with a frequency response high cut-off at 50 Hz (90 dB–octave roll off) and digitized at a rate of 250 samples per second. Digitized signals were recorded to disk using Brain Vision Recorder software (Brain Products GmbH, Munich, Germany). Interelectrode impedances were maintained below 20 kΩ. Two electrodes were also placed on the left and right mastoids. The EEG was recorded using the average reference. The electroocculogram (EOG) was recorded for the purpose of artifact correction; horizontal EOG was recorded from the external canthi of both eyes, and vertical EOG was recorded from the suborbit of the right eye and electrode channel Fp2.

### Data Analysis

#### Behavior

To recap, participants’ estimation of 1 second was considered on-time if it occurred within a narrow window of time centered around 1 second, the size of which was initialized at 200 ms, such that responses occurring between 900 and 1100 ms were considered correct at the start of the experiment. The time-window size was adjusted from trial to trial according to a staircase procedure that ensured 50% probabilities of receiving positive and negative feedback (see Task Design). The mean time-window size averaged across trials for each block was calculated for each participant. Therefore, smaller and larger average time-window sizes indicate, respectively, better and worse performance.

#### Electrophysiology

Post-processing and data visualization were performed using Brain Vision Analyzer software (Brain Products GmbH). The digitized signals were filtered using a fourth-order digital Butterworth filter with a passband of 0.10–30 Hz. The data were segmented for an 800 ms epoch extending from 200 ms prior to 600 ms following presentation of reward and no-reward feedback. Ocular artifacts were corrected using an eye movement correction algorithm (Gratton et al., 1983). The EEG data were re-referenced to averaged mastoids electrodes. Data were baseline corrected by subtracting from each sample for each channel the mean voltage associated with that electrode during the 200 ms interval preceding feedback onset. Trials with muscular and other artifacts were excluded according to a 150 μV Max-Min voltage difference, a ±150 μV level threshold, a ±35 μV step threshold, and a 0.1 μV lowest-allowed activity level as rejection criteria. ERPs were then created for each electrode and participant by averaging the single-trial EEG according to the reward and no-reward feedback type.

The RewP was measured at channel FCz, where it reached maximum amplitude, utilizing a difference wave approach that isolated the RewP from overlapping ERP components such as the P300 (Holroyd and Krigolson, 2007; Sambrook and Goslin, 2015); a difference wave was created for each participant by subtracting the ERP to no-reward feedback stimuli from the ERP to reward feedback stimuli. RewP amplitude was then determined by averaging the mean amplitude in the difference wave from 200 to 300 ms following feedback onset (determined based on a visual inspection of the grand-average of the difference wave across all participants, which ensured that the grand-average RewP in this time range was characterized by a positive peak with a frontal-central voltage distribution). For the purpose of comparison, P300 amplitude was measured at channel Pz, where it typically reaches maximum amplitude across the scalp (Donchin and Coles, 1988). Individual subject ERPs were averaged across feedback conditions, and P300 amplitude was defined by finding the maximum positive deflection from 280 to 420 ms following feedback onset (as determined based on a visual inspection of the grand-average ERP across all trials and all participants)^2^. In order to assess FMT power, the continuous EEG data were segmented into consecutive 4 seconds epochs with 200 ms overlap between segments (starting from the beginning of the experiment and continuing to the end) averaged across feedback types^3^. Artifact rejection and ocular correction were conducted on these EEG epochs as for the ERP data, and then submitted to a power spectral analysis using a Fast Fourier Transform (FFT) (Hanning Window, 10% length). FMT was assessed for each trial by averaging power between 4 and 8 Hz at each channel.

We employed a multi-level modeling analysis using the MIXED function in SPSS (IBM SPSS 24) in order to examine within-person changes in behavioral (i.e., time-window size) and electrophysiological measures (i.e., RewP, FMT) across the 16 blocks. To assess how these measures fluctuated across time, we calculated person-centered scores such that mean performance averaged across all blocks of trials of a given participant was subtracted from the mean performance averaged across trials for each block (e.g., adjusted time-window size for block 1 = average time-window size for block 1 – the average time-window size across all 16 blocks) (e.g., Saunders et al., 2015). By doing this, each participant’s score for a given measurement was calculated relative to their own average score, with positive and negative values indicating higher and lower values than average for that participant, respectively. We investigated the relation among within-person fluctuations in time-window size, RewP amplitude, and FMT power with time on task, with the time (block) variable coded both as a linear slope from 0 to 15 (reflecting the 16 blocks in total) and as a quadratic slope, as calculated as the square of the linear-time variable (i.e., 0^2^ to 15^2^). In total we tested three models, each separately testing the time-window size, RewP amplitude, and FMT power as a function of the other two variables and both the linear and quadratic time variables. For example, a model testing the time-course of RewP amplitude included time-window size, FMT power, linear-time, and quadratic-time as its predictors. All of the analyses were conducted with a random intercept for each participant, unstructured covariance type, and maximum likelihood estimation. Both time (block) variables were treated as random slopes. Effect size, *r*, is reported for each model effect.

For completeness, we also applied multi-level modeling analyses to examine between-subjects relationships among time-window size, RewP amplitude, and FMT power. Each participant’s average time-window size, RewP amplitude, and FMT power values were calculated separately across all the trials. Each score was then grand mean-centered for each participant by subtracting the grand average (across all participants) from each measure (e.g., the grand mean-centered RewP amplitude for a participant = the mean RewP amplitude averaged across all trials for this participant – the grand mean RewP amplitude averaged across all participants). In total we tested three models that separately tested time-window size, RewP amplitude, and FMT power as a function of the other two variables, each using a random intercept for each participant, unstructured covariance type, and maximum likelihood estimation.

We also explored the effects of personality traits using multi-level modeling analyses, the results of which are reported in the supplementary materials.

## Results

Three participants discontinued the task^4^. The data of three other participants were excluded from analysis due to self-reported neurological or psychiatric disorders. Further, during the first few blocks of trials one participant produced extreme time estimates (i.e., by responding about 7 seconds after cue onset); performance for this participant improved thereafter, but the probability of reward was strongly biased across blocks of trials due to the staircase procedure (mostly no reward in the first few blocks, followed by mostly reward in the following few blocks). The data of two participants were also excluded due to artifacts associated with excessive head movement. Further, we excluded data from three participants with average time-window size or FMT power that was 3 standard deviations (SD) above the group means. In total these exclusions resulted in the data of 53 participants used for all of the analyses.

### Behavior

#### Time-window size

Figure 1a depicts the time-course of the mean time-window size averaged across participants. The multi-level modeling analysis predicting the time-window size with RewP, FMT power, linear-time, and quadratic-time revealed significant main effects of linear-time and quadratic-time (Table 1, lines 1 & 2). These results confirm the impression of Figure 1a that participants’ performance improved with time but worsened towards the end of the experiment. We also found significant interactions of linear- and quadratic-time with FMT power on time-window size (Table 1, lines 5 & 7). Figure 2 reveals the nature of these effects. The relation between FMT power and the time-window size differed across time, such that greater FMT power (relative to participants’ own average FMT power) was associated with better (smaller) time-window size in the beginning of the experiment, as well as with relatively more stable performance with time on task (i.e., a relatively shallower change in performance over time). A significant main effect of FMT power (Table 1, line 3) signifies that the difference in time-window size by FMT power in the beginning was statistically significant.

**Figure 1.**
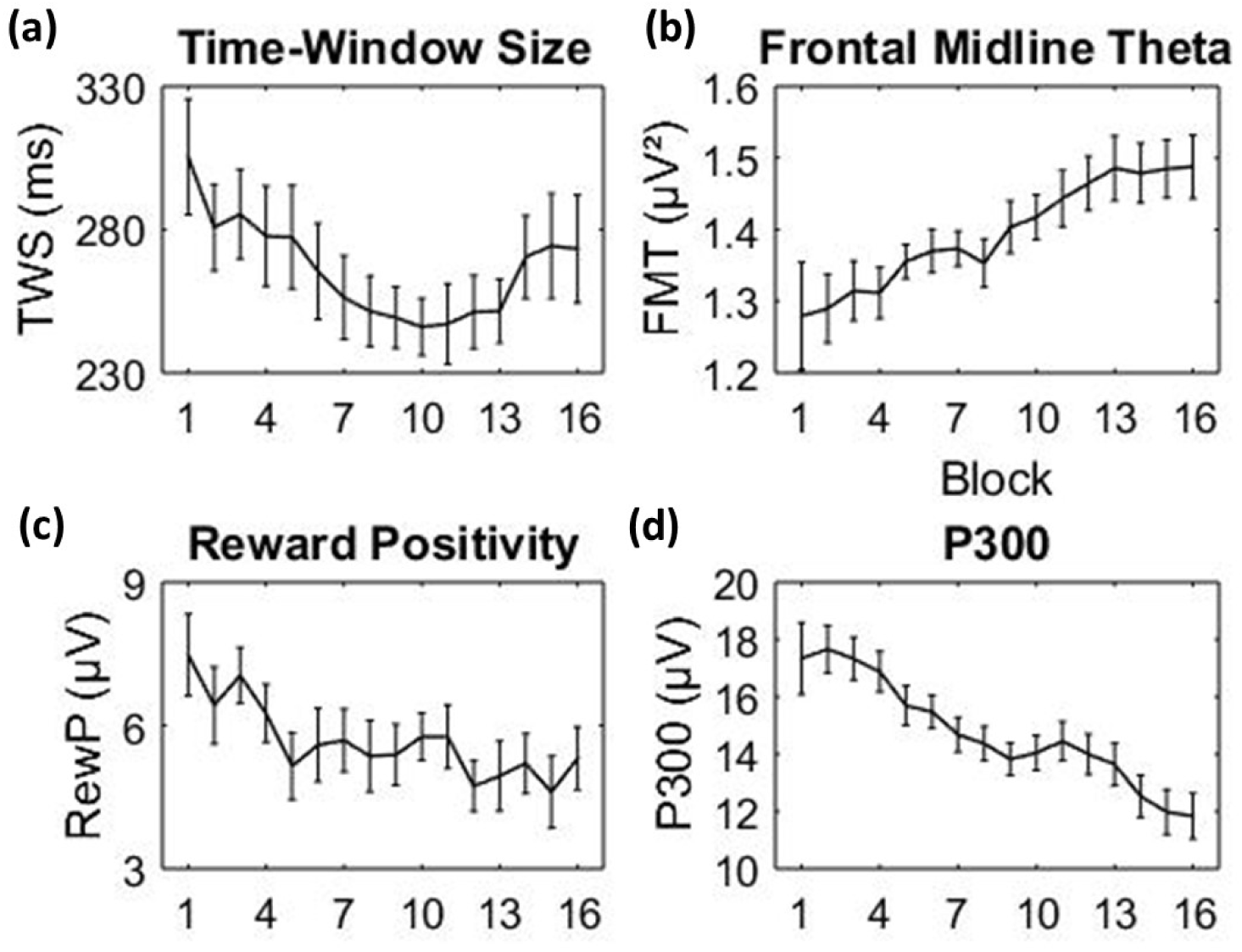
Block by block performance and electrophysiological measures, averaged across subjects for (a) time-window size, (b) frontal midline theta power, (c) reward positivity amplitude, and (d) P300 amplitude. Error bars indicate within-subject 95% confidence intervals (Cousineau, 2005).

**Figure 2.**
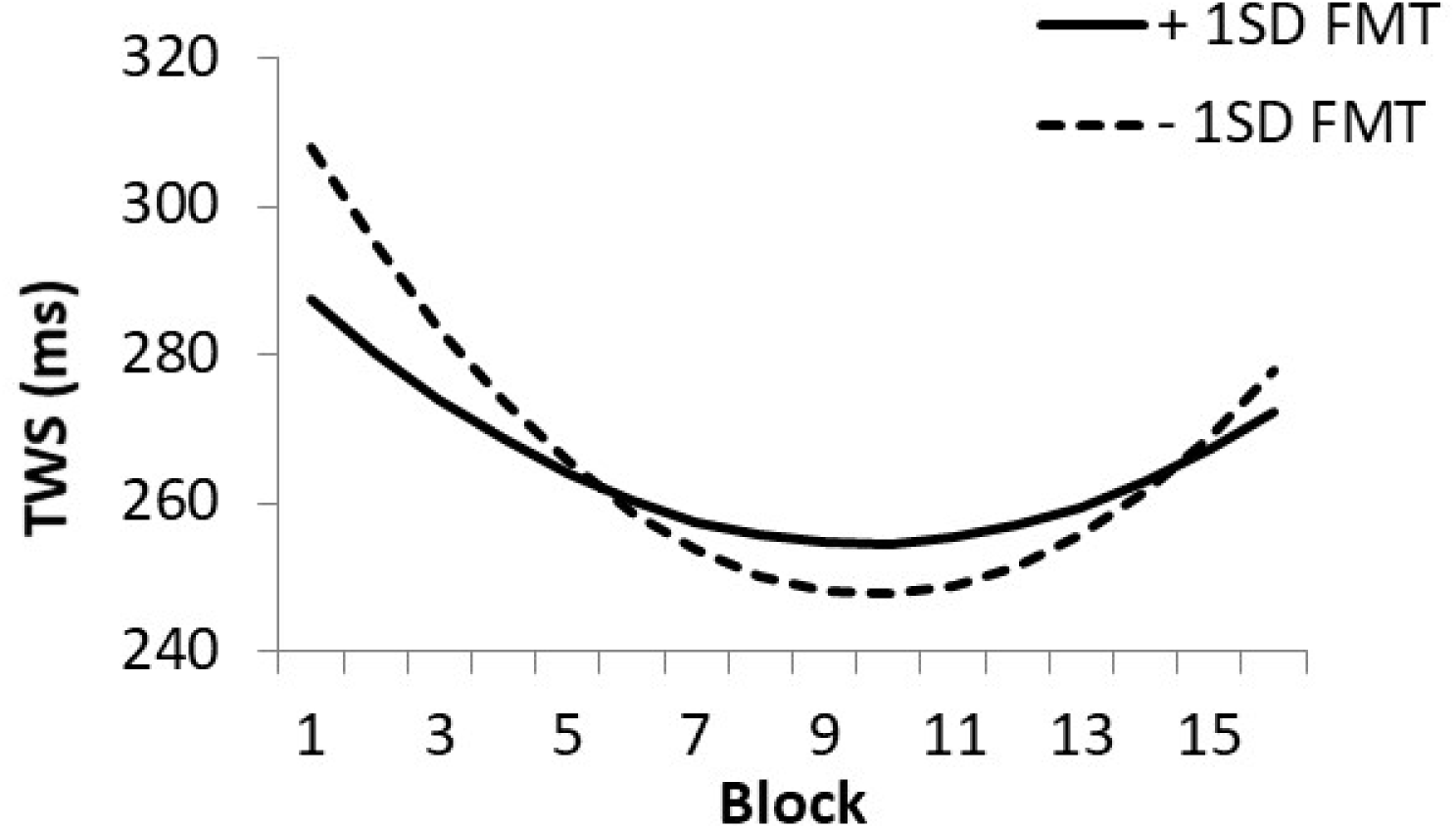
The within-subjects effect depicting the interaction of frontal midline theta (FMT) power with linear-time (block) and quadratic-time on time-window size (TWS). The solid and dotted lines denote TWS across blocks when participants produced FMT power that was above or below one standard deviation relative to their own average FMT, respectively (+/-1 SD FMT). Note that this figure is derived from the model predicting TWS based on time and FMT power (i.e., excluding reward positivity amplitude).

**Table 1.** Multi-level modeling results on time-window size (TWS), frontal midline theta (FMT) power, reward positivity (RewP) amplitude, and P300 amplitude with time-on-task. Line indicates the row number in which each effect is reported (for the purpose of indexing in the text). Block-L = block (time) coded as a linear slope from 0 to 15. Block-Q = block (time) coded as a quadratic slope (as the power of linear slope) from 0^2^ to 15^2^. Significant results (p<0.05) are highlighted in bold.

| DV | Predictors | b | SE | 95% CIs | r | pVal | Line |
| --- | --- | --- | --- | --- | --- | --- | --- |
| TWS | **Block-L** | **-10.28** | **2.41** | **[-15.1, -5.47]** | **0.478** | **<.001** | 1 |
|  | **Block-Q** | **0.6** | **0.16** | **[.27, .93]** | **0.424** | **0.001** | 2 |
|  | **FMT** | **-78.3** | **30.22** | **[-137.9, -18.71]** | **0.181** | **0.01** | 3 |
|  | RewP | 2.04 | 1.82 | [-1.53, 5.62] | 0.043 | 0.263 | 4 |
|  | **Block-L\*FMT** | **22.64** | **8.89** | **[5.2, 40.09]** | **0.093** | **0.011** | 5 |
|  | Block-L*RewP | -0.9 | 0.57 | [-2.02, .22] | 0.057 | 0.115 | 6 |
|  | **Block-Q\*FMT** | **-1.25** | **0.61** | **[-2.45, -.05]** | **0.074** | **0.041** | 7 |
|  | Block-Q*RewP | 0.06 | 0.04 | [-.01, .14] | 0.061 | 0.093 | 8 |
| FMT | **Block-L** | **0.01** | **0.003** | **[.01, .02]** | **0.532** | **<.001** | 9 |
|  | **TWS** | **-0.0003** | **0.0001** | **[-.0006, -.0001]** | **0.084** | **0.019** | 10 |
|  | RewP | 0.004 | 0.003 | [-.001, .01] | 0.053 | 0.142 | 11 |
|  | **Block-L\*TWS** | **0.00004** | **0.00002** | **[.00001, .00007]** | **0.091** | **0.012** | 12 |
|  | Block-L*RewP | -0.0004 | 0.0004 | [-.001, .0003] | 0.037 | 0.304 | 13 |
| RewP | **Block-L** | **-0.35** | **0.09** | **[-.53, -.17]** | **0.453** | **<.001** | 14 |
|  | **Block-Q** | **0.01** | **0.01** | **[.004, .02]** | **0.326** | **0.009** | 15 |
|  | TWS | 0.007 | 0.004 | [-.001, .01] | 0.070 | 0.087 | 16 |
|  | FMT | 1.27 | 1.46 | [-1.61, 4.16] | 0.064 | 0.385 | 17 |
|  | **Block-L\*TWS** | **-0.003** | **0.001** | **[-.006, -.0006]** | **0.087** | **0.015** | 18 |
|  | Block-L*FMT | 0.05 | 0.45 | [-.83, .94] | 0.004 | 0.909 | 19 |
|  | **Block-Q\*TWS** | **0.0002** | **0.00008** | **[.00004, .0004]** | **0.087** | **0.014** | 20 |
|  | Block-Q*FMT | -0.01 | 0.03 | [-.07, .05] | 0.013 | 0.724 | 21 |
| P300 | **Block-L** | **-0.38** | **0.05** | **[-.47, -.29]** | **0.755** | **<.001** | 22 |

### Electrophysiology

#### Frontal Midline Theta

Figure 1b depicts the time-course of FMT power over time averaged across participants. As the inclusion of a quadratic time variable in the model did not reveal a significant effect, the multi-level modeling analysis here tested a model excluding this variable. The model predicting FMT power with time-window size, RewP amplitude, and linear-time revealed a significant main effect of linear-time, such that FMT power increased over time (Table 1, line 9). There was also a significant interaction of linear-time and time-window size on FMT power (Table 1, line 12; see Supplementary Figure 1). The pattern of interaction indicates that better task performance was associated with larger FMT power at the beginning (as shown by the significant main effect of time-window size; Table 1, line 10) and less of an increase in FMT power with time on task (Supplementary Figure 1).

#### RewP

Figure 1c depicts the time-course of RewP amplitude averaged across participants, and Figure 3 depicts the RewP waveforms averaged across participants for every half-hour of task performance. The multi-level modeling analysis predicting RewP amplitude with time-window size, FMT power, linear-time, and quadratic-time revealed statistically significant linear (Table 1, line 14) and quadratic effects (Table 1, line 15) of time on RewP amplitude. This result indicates a nonlinear reduction in RewP amplitude (more negative) with time (Figure 1c). There were also significant interactions of linear-time and time-window size (Table 1, line 18) and of quadratic-time and time-window size (Table 1, line 20) on RewP amplitude, such that smaller (better) time-window size (relative to each participant’s own average) was associated with a relatively smaller, linear decline in RewP amplitude (Figure 4).

**Figure 3.**
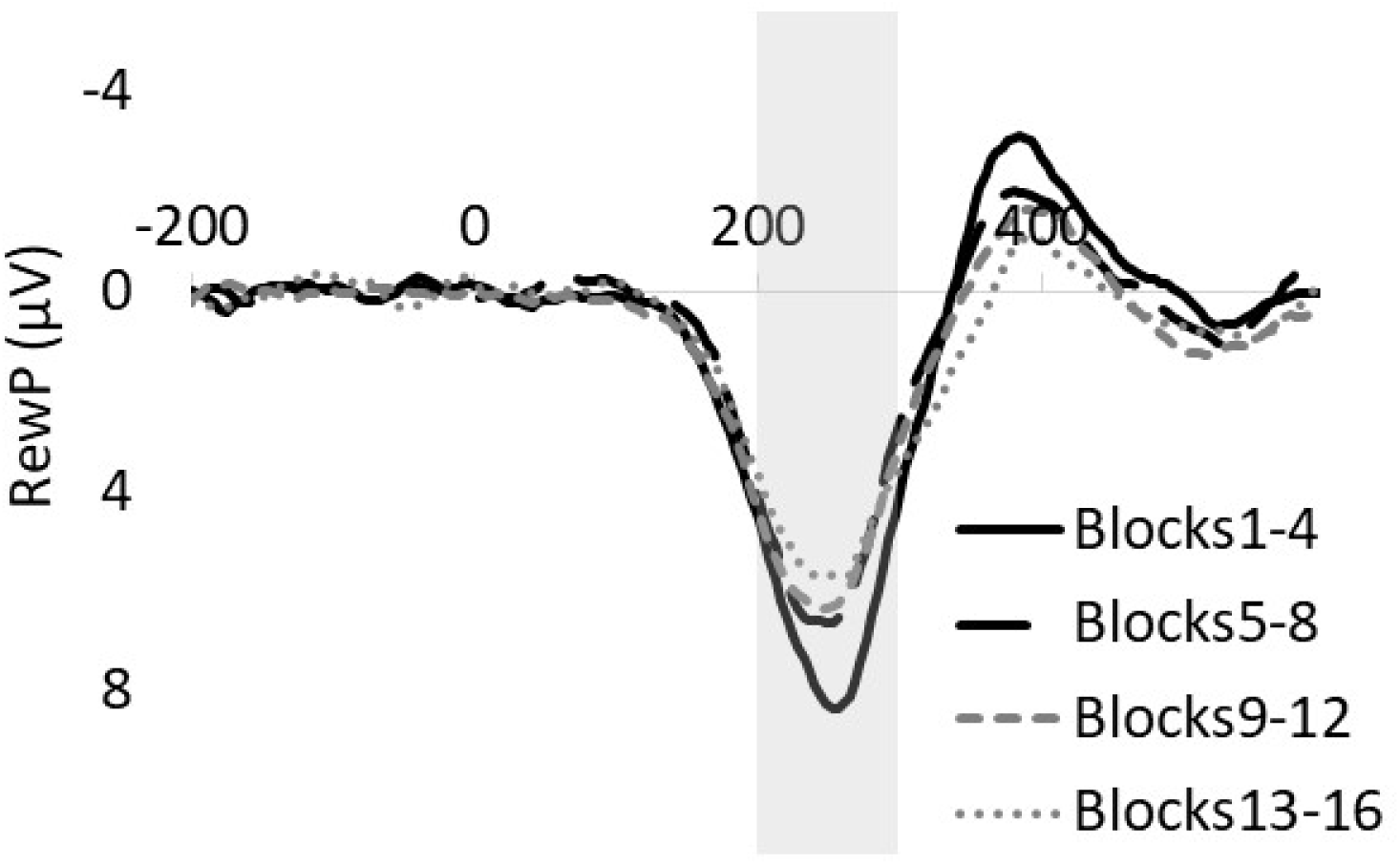
The reward positivity (RewP) derived as a difference wave (reward feedback minus no-reward feedback) to event-related potentials averaged across participants and across blocks, separately for every consecutive four blocks. Blocks 1-4 = RewP difference wave for block 1 to 4 (i.e., 1^st^ 30 min). Blocks 5-8 = RewP difference wave for block 5 to 8 (2^nd^ 30 min). Blocks 9-12 = RewP difference wave for block 9 to 12 (3^rd^ 30 min). Blocks 13-16 = RewP difference wave for block 13 to 16 (4^th^ 30 min). RewP amplitude was measured between 200-300 ms following feedback onset at time 0 (as highlighted in gray). Negative is plotted up by convention.

**Figure 4.**
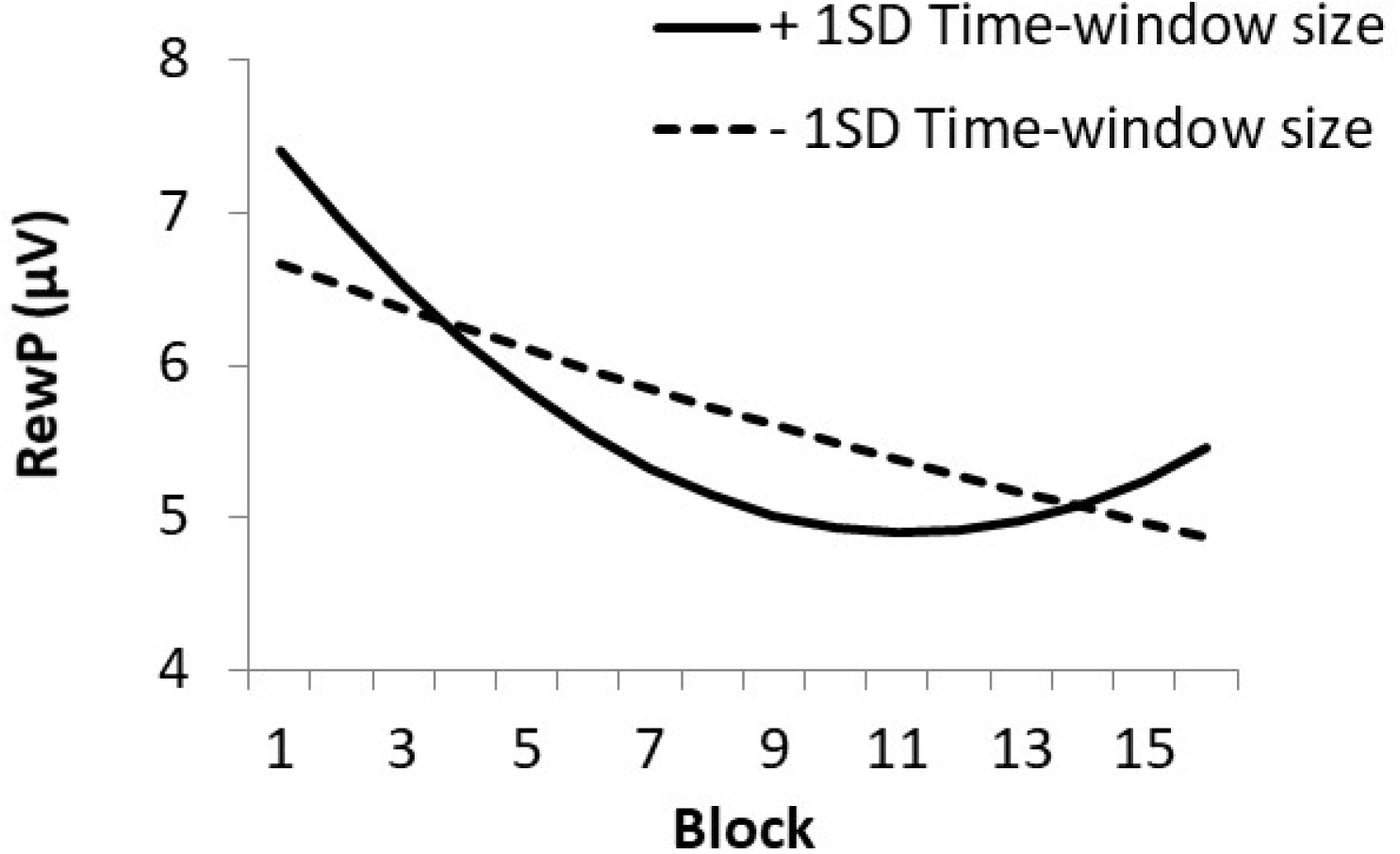
The within-subjects effect depicting the interaction of time-window size with linear-time (block) and quadratic-time on reward positivity (RewP) amplitude. The solid and dotted lines denote RewP amplitude across blocks when each participant’s time-window size was one standard deviation above or below their own average time-window size, respectively (+/-1 SD time-window size). Note that this figure is derived from the model predicting RewP amplitude based on linear-time, quadratic-time, and time-window size (excluding FMT power).

#### P300

Figure 1d depicts the time-course of P300 amplitude over time averaged across participants. As the inclusion of a quadratic time variable in the model did not reveal a significant effect, the multi-level modeling analysis here tested a model excluding this variable. A multi-level modeling analysis predicting P300 with linear-time revealed a significant main effect of linear-time (Table 1, line 22), indicating that P300 amplitude decreased linearly with time on task.

### Relationship between behavioral and electrophysiological measures

The multi-level modeling analysis examining the between-subjects relationships among time-window size, RewP, and FMT power did not reveal any statistically significant relationships (p>.40). This indicates that the overall sizes of FMT power and RewP amplitude were not related to task performance nor to each other. Note that because the statistical power for the between-participants analyses is low, these results should be interpreted with caution (as is also the case for the analyses related to personality, below).

### Personality questionnaires

A summary of each questionnaire score is provided in Table 2, and zero-order correlations among questionnaires are provided in Table 3.

We conducted a simple correlation analysis among time-window size, RewP, FMT, personality scores, and the task engagement level assessed by the post-experiment paper-and-pencil questionnaires (see Task Procedure)^5^. We observed that higher levels of task engagement were correlated with higher reward responsiveness scores (Pearson r=.310, p=.03), higher persistence scores (r=.316, p=.027), as well as better (smaller) TWS (r=-.381, p=.007). Exploratory analyses examining the impact of personality traits on time-window size, RewP amplitude, and FMT power did not reveal any significant effects of personality on these measures (see Supplementary Table 1).

**Table 2.** Summary of participant questionnaire scores. Temporal Experience of Pleasure-C= consummatory pleasure subscale of the temporal experience of pleasure scale. Temporal Experience of Pleasure-A= anticipatory pleasure subscale of the temporal experience of pleasure scale. Apathy is based on the apathy evaluation scale. Depression is based on the depression subscale of the depression, anxiety, and stress scale (DASS-21).

|  | *Mean* | *SD* | *Range* |
| --- | --- | --- | --- |
| Reward Responsiveness | 26.4 | 2.57 | 19-30 |
| Temporal Experience of Pleasure-C | 37.3 | 4.44 | 26-46 |
| Temporal Experience of Pleasure-A | 44.62 | 6.24 | 32-60 |
| Persistence | 67.91 | 10.49 | 42-87 |
| Apathy | 31.83 | 5.41 | 23-43 |
| Depression | 3.96 | 4.34 | 0-18 |

**Table 3.** Zero-order correlations among questionnaire scores.

|  | **1** | **2** | **3** | **4** | **5** |
| --- | --- | --- | --- | --- | --- |
| 1. Reward Responsiveness |  |  |  |  |  |
| 2. Temporal Experience of Pleasure-C | .03 |  |  |  |  |
| 3. Temporal Experience of Pleasure-A | .44** | .20 |  |  |  |
| 4. Persistence | .49** | -.04 | .26 |  |  |
| 5. Apathy | -.62** | -.17 | -.54** | -.58** |  |
| 6. Depression | -.33* | .04 | -.28* | -.57** | .40** |

\* $p < .05$ , \*\* $p < .01$

## Discussion

Goal-directed behaviors often require the deployment of cognitive control for their successful execution. Yet, because cognitive control is perceived to be effortful, people tend to avoid applying it. Theories of control have therefore proposed the existence of a neurocognitive mechanism that regulates control levels by weighing the benefits (or rewards) of control against its associated effort-related costs (Botvinick and Braver, 2015; Kurzban et al., 2013; Inzlicht et al., 2014; Shenhav et al., 2013, 2017; Westbrook and Braver, 2015). How this occurs is not fully understood, but ACC is believed to be partly responsible for it (e.g., Holroyd and Umemoto, 2016; Shenhav et al., 2013, 2017; Vassena et al., 2017; Verguts et al., 2015). Here we investigated the role of ACC in valuating and regulating control levels in order to sustain performance on an extended task, as revealed by electrophysiological indices of ACC activity (Holroyd and Yeung, 2012; Holroyd and McClure, 2015; Holroyd and Umemoto, 2016).

Participants performed a standard time estimation task for 2 hours while their EEG was recorded (Miltner et al., 1997). They received a small monetary reward for each “on-time” estimate, the difficulty of which was adjusted across trials so that all of the participants received the rewards on approximately 50% of the trials. Moreover, in order to increase their motivation the participants were told, truthfully, that better performance increased their chances for receiving one of two $100 Amazon gift cards, which in fact were later awarded to two of the participants via a lottery. We observed that time-window size decreased over time, following both a linear and a quadratic trend (Figure 1a); that is, performance initially improved with time-on-task and later deteriorated towards the end of the experiment. A straightforward interpretation of these findings is that the initial improvement in task performance reflects an early learning process, and that the later decrement in performance reflects an impairment due to mental fatigue. Consistent with this interpretation, post-experiment self-reports revealed that better (smaller) time-window size correlated with overall task engagement, suggesting that task performance depended strongly on the control levels applied; thus, a withdrawal of control should result in impaired performance. Further, we also observed that the amplitudes of the RewP and P300 decreased with time on task (Figure 1c & 1d), whereas FMT power increased with time on task (Figure 1b), as observed previously (Barwick et al., 2012; Paus et al., 1997; Wascher et al., 2014). These findings suggest that reward valuation and attention levels decreased while control levels increased with time-on-task. Taken together, they appear to indicate that greater application of control in order to counteract deteriorating levels of attention and reward valuation is associated with accumulating mental fatigue, as suggested by a deterioration in performance at the end of the task (see also Hopstaken, et al., 2015; c.f. Milyavskaya et al., 2017, in this issue).

Notably, the between-person analyses did not reveal any significant associations among time-window size, RewP amplitude, and FMT power, even though both of the electrophysiological phenomena are believed to be produced by ACC (Holroyd and Umemoto, 2016). Although low statistical power may have obscured these relationships, their absence may also suggest that these measures depend more strongly on within-participant differences than on between-participant differences. On this view, because reward valuation and effort expenditure provide semi-independent determinants of task performance, RewP amplitude and FMT power might not be expected to co-vary together across participants. Instead, within-subject measures might provide greater insight into these factors, as in fact we observed.

To wit, the within-person analysis predicting time-window size revealed significant linear and quadratic effects of time-on-task, and this effect was modulated by FMT power (Figure 2): greater FMT power (relative to each participant’s own average power) was associated with better (smaller) time-window sizes initially, and with relatively shallower changes in performance across time afterward. These findings provide insight into the role of control in regulating task performance: Although increased FMT power is associated with deteriorating performance at the end of the task, as discussed above (see Figures 1a and 1b), the within-subjects analysis reveals that greater FMT power is also associated with better performance at the start of the task (Figure 2). Greater FMT power is further associated with less steep of a fall in performance toward the end of the task. Thus, FMT power appears to reflect a neural process that enhances performance, both during an early stage when subjects learn to perform the task better, and a later stage when performance suffers due to cognitive fatigue.

Within-person analyses predicting RewP amplitude also revealed significant interactions of time-window size with both linear- and quadratic-time trends. That is, smaller (better) time-window size (relative to each participant’s own average time-window size) was associated with smaller declines in RewP amplitude over time, whereas larger (worse) time-window size was associated with a steeper decline in RewP amplitude with time on task (Figure 4). Similarly, within-person analyses predicting FMT revealed a significant interaction of linear-time and time-window size: Better task performance was associated with larger FMT power at the beginning of the task and smaller increases in FMT power with time-on-task (Supplementary Figure 1). It appears that greater task engagement, as reflected in smaller time-window sizes, was associated with more gradual reductions in RewP amplitude and more gradual increases in FMT power. That is, good task performance was associated with more stable indicators of reward valuation and cognitive control throughout the task.

Finally, we found from post-experiment self-reports that higher levels of task engagement correlated significantly with higher reward responsiveness and persistence scores, indicating that participants who scored high on these personality traits were more engaged during the entire experiment. An exploration of whether persistence and reward sensitivity were associated with task performance, RewP amplitude, and FMT power failed to reveal any significant relations among them (see Supplementary Table 1). Although a number of studies have reported individual differences in personality associated with RewP amplitude (e.g., Bress and Hajcak, 2013, Cooper et al., 2014; Liu et al., 2014; Parvaz et al., 2016; Umemoto and Holroyd, 2017) and phasic FMT power (e.g., Cavanagh and Shackman, 2014), we did not replicate any of these findings. Nevertheless, such effects may have been obscured by the low statistical power in the present study for conducting analyses related to individual differences. Adequate sample sizes will be needed in future studies to elucidate the role of personality in extended task performance (e.g., Fraley and Vazire, 2014; see also Umemoto and Holroyd, 2016) and its expression in individual differences in electrophysiological correlates of ACC function (Holroyd and Umemoto, 2016).

A challenging question is whether increasing FMT power with time-on-task reflects growing mental fatigue per se, or the application of effortful control needed to overcome that fatigue, as these processes are necessarily correlated. In fact, several neuroimaging studies have implicated ACC in cognitive fatigue (e.g., Cook et al., 2007; Dobryakova et al., 2013; Wylie et al., 2017). However, we argue against the simple association between FMT and fatigue itself. Specifically, mental fatigue is generally associated with impaired task performance (as reflected by increased error rates and greater reaction times), but performance here improved over the first 90 min of the study while FMT power continued to rise. Furthermore, during the initial stage of the task greater FMT was associated with better task performance. Our observation that changes in FMT power closely followed time-on-task performance, as well as relatively more stable (shallower changes in) task performance, appears inconsistent with the account that FMT simply reflects mental fatigue itself.

Past studies have shown that a brief, phasic burst of FMT power is commonly elicited by response conflict and error commission (e.g., Cavanagh and Frank, 2014). The amplitude of the error related negativity (ERN), a performance-related ERP component associated with both the RewP (Holroyd and Coles, 2002) and FMT power (Cavanagh and Frank, 2014), has been observed to decline with time on task (Lorist et al 2005; Boksem et al 2006; Inzlicht and Gutsell, 2007). By contrast, ongoing FMT power, as we measured in this study, is associated with prolonged mental challenges and high working memory load (Hsieh and Ranganath, 2014; Mitchell et al., 2008). Understanding how phasic and ongoing FMT signals relate to each other is an important question for future studies. For example, past studies have shown that the adverse effects of mental fatigue on task performance and ERN amplitude, which is associated with phasic FMT (Cavanagh and Frank, 2014), can be counteracted with motivational incentives (e.g., Boksem et al., 2006; Hopstaken et al., 2015; Sarter et al., 2006). Whether fatigue-related changes in tonic FMT power can also be reversed with task incentives is an interesting avenue for future research.

In summary, although task performance improved until the last quarter of the experiment, when it started to reverse, decreasing RewP and P300 amplitudes indicated that the participants gradually disengaged from the task and devalued the outcomes of their performance. By contrast, FMT power increased monotonically with time on task, suggesting greater recruitment of cognitive control even as the task became harder and less rewarding. Although task performance and the electrophysiological measures did not correlate with each other across participants, a within-person increase in FMT power was associated with 1) better task performance at the start of the experiment, suggesting that high control levels facilitated learning; 2) more stable performance overall. Conversely, better task performance was associated with higher self-reports of task engagement and relatively more stable RewP amplitudes and FMT power over time. Together, we interpret these results as reflecting the role of ACC in sustaining behavior over an extended period, especially for tasks that demand high levels of cognitive control with low immediate reward value.

## Acknowledgements

This work was supported by a grant from the Japan Society for the Promotion of Science (A.U.) and Natural Sciences and Engineering Research Council of Canada Discovery Grant 312409-05 (C.B.H.).

## Footnotes

1 The results of these analyses are provided in the supplementary materials. Note that this sample size only provides adequate power to detect medium-to-large effects related to individual differences in personality.

2 Note that the scalp distribution for P300 was not maximal centrally but rather over lateral scalp areas (e.g., electrodes P3 and P4). However, given that the P300 was not the focus of our analysis, P300 amplitude was measured at channel Pz in keeping with previous studies, and because P300 amplitude was maximal there when comparing only the midline channels.

3 When we segmented the EEG data with a 50% overlap between segments for the frontal midline theta analyses, there was no significant difference in the overall power (p=0.415), and all the results remained the same.

4 One participant indicated signs of claustrophobia, one participant left feeling unwell, and one participant discontinued after reporting that s/he had earned enough money.

5 Note that four of the participants’ engagement levels were not obtained due to an error, hence the number of participants included in this analysis was 49.

## References

Asada, H., Fukuda, Y., Tsunoda, S., Yamaguchi, M., Tonoike, M., 1999. Frontal midline theta rhythms reflect alternative activation of prefrontal cortex and anterior cingulate cortex in humans. Neurosci. Lett. 274, 29–32. DOI:10.1016/S0304-3940(99)00679-5

Barwick, F., Arnett, P., Slobounov, S., 2012. EEG correlates of fatigue during administration of a neuropsychological test battery. Clin. Neurophysiol. 123, 278–284. DOI:10.1016/j.clinph.2011.06.027

Becker, M.P.I., Nitsch, A.M., Miltner, W.H.R., Straube, T., 2014. A single-trial estimation of the feedback-related negativity and its relation to BOLD responses in a time-estimation task. J. Neurosci. 34, 3005–12. DOI:10.1523/JNEUROSCI.3684-13.2014

Boksem, M.A.S., Meijman, T.F., Lorist, M.M., 2006. Mental fatigue, motivation and action monitoring. Biol. Psychol. 72, 123–132. DOI:10.1016/j.biopsycho.2005.08.007

Boksem, M.A.S., Tops, M., 2008. Mental fatigue: Costs and benefits. Brain Res. Rev. 59, 125– 139. DOI:10.1016/j.brainresrev.2008.07.001

Botvinick, M., Braver, T., 2015. Motivation and Cognitive Control: From Behavior to Neural Mechanism. Annu. Rev. Psychol. 66, 83–113. DOI:10.1146/annurev-psych-010814-015044

Botvinick, M.M., 2012. Hierarchical reinforcement learning and decision making. Curr. Opin.Neurobiol. 22, 956–962. DOI:10.1016/j.conb.2012.05.008

Botvinick, M.M., Niv, Y., Barto, A.C., 2009. Hierarchically organized behavior and its neural foundations: A reinforcement learning perspective. Cognition 113, 262–280. DOI:10.1016/j.cognition.2008.08.011

Brainard, D.H., 1997. The Psychophysics Toolbox. Spat. Vis. 10, 433–436. DOI:10.1163/156856897X00357

Bress, J.N., Hajcak, G., 2013. Self-report and behavioral measures of reward sensitivity predict the feedback negativity. Psychophysiology 50, 610–616. DOI:10.1111/psyp.12053

Cavanagh, J.F., Frank, M.J., 2014. Frontal theta as a mechanism for cognitive control. Trends Cogn. Sci. 18, 414–421. DOI:10.1016/j.tics.2014.04.012

Cavanagh, J.F., Shackman, A.J., 2014. Frontal midline theta reflects anxiety and cognitive control: Meta-analytic evidence. J. Physiol. Paris 109, 3–15.DOI:10.1016/j.jphysparis.2014.04.003

Cloninger, C.R., Svrakic, D.M., Przybeck, T.R., 1993. A psychobiological model of temperament and character. Arch. Gen. Psychiatry 50, 975–990.

Cook, D.B., O’Connor, P.J., Lange, G., Steffener, J., 2007. Functional neuroimaging correlates of mental fatigue induced by cognition among chronic fatigue syndrome patients and controls. Neuroimage 36, 108–122. DOI:10.1016/j.neuroimage.2007.02.033

Cooper, A.J., Duke, E., Pickering, A.D., Smillie, L.D., 2014. Individual differences in reward prediction error: contrasting relations between feedback-related negativity and trait measures of reward sensitivity, impulsivity and extraversion. Front. Hum. Neurosci. 8, 248. DOI:10.3389/fnhum.2014.00248

Dobryakova, E., DeLuca, J., Genova, H.M., Wylie, G.R., 2013. Neural Correlates of Cognitive Fatigue: Cortico-Striatal Circuitry and Effort–Reward Imbalance. J. Int. Neuropsychol. Soc. 19, 849–853. DOI:10.1017/S1355617713000684

Donchin, E., Coles, M.G.H., 1988. Is the P300 component a manifestation of context updating? Behav. Brain Sci. 11, 357–374. DOI:10.1017/S0140525X00058015

Ebitz, R.B., Hayden, B.Y., 2016. Dorsal anterior cingulate: a Rorschach test for cognitive neuroscience. Nat. Neurosci. 19, 1278–1279. DOI:10.1038/nn.4387

Erdfelder, E., Faul, F., Buchner, A., 1996. GPOWER: A general power analysis program. Behav. Res. Methods, Instruments, Comput. 28, 1–11. DOI:10.3758/BF03203630

Fraley, R.C., Vazire, S., 2014. The N-pact factor: Evaluating the quality of empirical journals with respect to sample size and statistical power. PLoS One 9. DOI:10.1371/journal.pone.0109019

Gard, D.E., Gard, M.G., Kring, A.M., John, O.P., 2006. Anticipatory and consummatory components of the experience of pleasure: A scale development study. J. Res. Pers. 40, 1086–1102. DOI:10.1016/j.jrp.2005.11.001

Gratton, G., Coles, M.G.H., Donchin, E., 1983. A new method for off-line removal of ocular artifact. Electroencephalogr. Clin. Neurophysiol. 55, 468–484. DOI:10.1016/0013-4694(83)90135-9

Gusnard, D. a, Ollinger, J.M., Shulman, G.L., Cloninger, C.R., Price, J.L., Van Essen, D.C., Raichle, M.E., 2003. Persistence and brain circuitry. Proc. Natl. Acad. Sci. U. S. A. 100, 3479–3484. DOI:10.1073/pnas.0538050100

Hockey, G.R.J., 2011. A motivational control theory of cognitive fatigue. Cogn. fatigue Multidiscip. Perspect. Curr. Res. Futur. Appl. 167–187. DOI:10.1037/12343-008

Holroyd, C.B., Coles, M.G.H., 2002. The Neural Basis of Human Error Processing: Reinforcement Learning, Dopamine, and the Error-Related Negativity. Psychol. Rev. 109, 679–709. DOI:10.1037//0033-295X.109.4.679

Holroyd, C.B., Krigolson, O.E., 2007. Reward prediction error signals associated with a modified time estimation task. Psychophysiology 44, 913–917. DOI:10.1111/j.1469-8986.2007.00561.x

Holroyd, C.B., McClure, S.M., 2015. Hierarchical Control Over Effortful Behavior by Rodent Medial Frontal Cortexu: A Computational Model. Psychol. Rev. 122, 54–83.

Holroyd, C.B., Umemoto, A., 2016. The research domain criteria framework: The case for anterior cingulate cortex. Neurosci. Biobehav. Rev. 71, 418–443. DOI:10.1016/j.neubiorev.2016.09.021

Holroyd, C.B., Yeung, N., 2012. Motivation of extended behaviors by anterior cingulate cortex. Trends Cogn. Sci. 16, 122–128. DOI:10.1016/j.tics.2011.12.008

Hopstaken, J.F., van der Linden, D., Bakker, A.B., Kompier, M.A.J., 2015. A multifaceted investigation of the link between mental fatigue and task disengagement. Psychophysiology 52, 305–315. DOI:10.1111/psyp.12339

Hsieh, L.T., Ranganath, C., 2014. Frontal midline theta oscillations during working memory maintenance and episodic encoding and retrieval. Neuroimage 85, 721–729. DOI:10.1016/j.neuroimage.2013.08.003

Ishii, R., Shinosaki, K., Ukai, S., Inouye, T., Ishihara, T., Yoshimine, T., Hirabuki, N., Asada, H., Kihara, T., Robinson, S.E., Takeda, M., 1999. Medial prefrontal cortex generates frontal midline theta rhythm. Neuroreport 10, 675–679. DOI:10.1097/00001756-199903170-00003

Inzlicht, M., Bartholow, B.D., Hirsh, J.B., 2015. Emotional foundations of cognitive control. Trends Cogn. Sci. 19, 126–132. DOI:10.1016/j.tics.2015.01.004

Inzlicht, M., Gutsell, J.N., 2007. Running on empty: Neural signals for self-control failure. Psychol. Sci. 18, 933–937. DOI:10.1111/j.1467-9280.2007.02004.x

Inzlicht, M., Schmeichel, B.J., Macrae, C.N., 2014. Why self-control seems (but may not be) limited. Trends Cogn. Sci. 18, 127–133. DOI:10.1016/j.tics.2013.12.009

Janssen, D.J.C., Poljac, E., Bekkering, H., 2016. Binary sensitivity of theta activity for gain and loss when monitoring parametric prediction errors. Soc. Cogn. Affect. Neurosci. 11, 1280–1289. DOI:10.1093/scan/nsw033

Jasper, H.H., 1958. The ten twenty electrode system of the international federation. Electroencephalogr. Clin. Neurophysiol. 10, 371–375.

Kool, W., McGuire, J.T., Rosen, Z.B., Botvinick, M.M., 2010. Decision making and the avoidance of cognitive demand. J. Exp. Psychol. Gen. 139, 665–682. DOI:10.1037/a0020198

Kurzban, R., Duckworth, A., Kable, J.W., Myers, J., 2013. An opportunity cost model of subjective effort and task performance. Behav. Brain Sci. 36, 661–679. DOI:10.1017/S0140525X12003196

Lin, H., Saunders, B., Hutcherson, C.A., Inzlicht, M (2017, August). Midfrontal theta and pupil dilation track subjective conflict in value-based decisions. Poster session presented at the 13th International Conference for Cognitice Neuroscience, Amsterdam, Netherlands.

Liu W. hua, Wang L. zhi, Shang H. rui, Shen, Y., Li, Z., Cheung, E.F.C., Chan, R.C.K., 2014. The influence of anhedonia on feedback negativity in major depressive disorder. Neuropsychologia 53, 213–220. DOI:10.1016/j.neuropsychologia.2013.11.023

Lorist, M.M., Boksem, M.A.S., Ridderinkhof, K.R., 2005. Impaired cognitive control and reduced cingulate activity during mental fatigue. Cogn. Brain Res. 24, 199–205. DOI:10.1016/j.cogbrainres.2005.01.018

Lovibond, P.F., Lovibond, S.H., 1995. The structure of negative emotional states: Comparison of the Depression Anxiety Stress Scales (DASS) with the Beck Depression and Anxiety Inventories. Behav. Res. Ther. 33, 335–343. DOI:10.1016/0005-7967(94)00075-U

Luu, P., Tucker, D.M., 2001. Regulating action: Alternating activation of midline frontal and motor cortical networks. Clin. Neurophysiol. 112, 1295–1306. DOI:10.1016/S1388-2457(01)00559-4

Marin, R.S., Biedrzycki, R.C., Firinciogullari, S., 1991. Reliability and validity of the apathy evaluation scale. Psychiatry Res. 38, 143–162. DOI:10.1016/0165-1781(91)90040-V

McGuire, J.T., Botvinick, M.M., 2010. Prefrontal cortex, cognitive control, and the registration of decision costs. Proc. Natl. Acad. Sci. 107, 7922–7926. DOI:10.1073/pnas.0910662107

Milyavskaya, M., Inzlicht, M., Johnson, T., Larson, M (submitted in this issue). Reward sensitivity following boredom and cognitive effort: A high-powered neurophysiological investigation.

Miltner, W.H.R., Braun, C.H., Coles, M.G.H., 1997. Event-related brain potentials following incorrect feedback in a time-estimation task: evidence for a “generic” neural system for error detection.

Mitchell, D.J., McNaughton, N., Flanagan, D., Kirk, I.J., 2008. Frontal-midline theta from the perspective of hippocampal “theta.” Prog. Neurobiol. 86, 156–185. DOI:10.1016/j.pneurobio.2008.09.005

Nelson, B.D., Kessel, E.M., Jackson, F., Hajcak, G., 2016. The impact of an unpredictable context and intolerance of uncertainty on the electrocortical response to monetary gains and losses. Cogn. Affect. Behav. Neurosci. 16, 153–163. DOI:10.3758/s13415-015-0382-3

Norman, D.A., Shallice, T., 1986. Attention to Action. Conscious. Self-Regulation 4, 1–18. DOI:10.1007/978-1-4757-0629-1_1

Parvaz, M.A., Gabbay, V., Malaker, P., Goldstein, R.Z., 2016. Objective and specific tracking of anhedonia via event-related potentials in individuals with cocaine use disorders. Drug Alcohol Depend. 164, 158–165. DOI:10.1016/j.drugalcdep.2016.05.004

Paus, T., Zatorre, R.J., Hofle, N., Caramanos, Z., Gotman, J., Petrides, M., Evans, A.C., 1997. Time-Related Changes in Neural Systems Underlying Attention and Arousal During the Performance of an Auditory Vigilance Task. J. Cogn. Neurosci. 9, 392–408. DOI:10.1162/jocn.1997.9.3.392

Pelli, D.G., 1997. The VideoToolbox software for visual psychophysics: transforming numbers into movies. Spat. Vis. DOI:10.1163/156856897X00366

Proudfit, G.H., 2015. The reward positivity: From basic research on reward to a biomarker for depression. Psychophysiology 52, 449–459. DOI:10.1111/psyp.12370

Sambrook, T.D., Goslin, J., 2015. A neural reward prediction error revealed by a meta-analysis of ERPs using great grand averages. Psychol. Bull. 141, 213–235. DOI:10.1037/bul0000006

Sarter, M., Gehring, W.J., Kozak, R., 2006. More attention must be paid: The neurobiology of attentional effort. Brain Res. Rev. 51, 145–160. DOI:10.1016/j.brainresrev.2005.11.002

Saunders, B., Milyavskaya, M., Inzlicht, M., 2015. What does cognitive control feel like? Effective and ineffective cognitive control is associated with divergent phenomenology. Psychophysiology 52, 1205–1217. DOI:10.1111/psyp.12454

Scheeringa, R., Bastiaansen, M.C.M., Petersson, K.M., Oostenveld, R., Norris, D.G., Hagoort, P., 2008. Frontal theta EEG activity correlates negatively with the default mode network in resting state. Int. J. Psychophysiol. 67, 242–251. DOI:10.1016/j.ijpsycho.2007.05.017

Shenhav, A., Botvinick, M.M., Cohen, J.D., 2013. The expected value of control: An integrative theory of anterior cingulate cortex function. Neuron 79, 217–240. DOI:10.1016/j.neuron.2013.07.007

Shenhav, A., Musslick, S., Lieder, F., Kool, W., Griffiths, T.L., Cohen, J.D., Botvinick, M.M., 2017. Toward a Rational and Mechanistic Account of Mental Effort. Annu. Rev. Neurosci. 40, 99–124. DOI:10.1146/annurev-neuro-072116-031526

Treynor, W., Gonzalez, R., Nolen-Hoeksema, S., 2003. Rumination reconsidered: A psychometric analysis. Cognit. Ther. Res. 27, 247–259.

Umemoto, A., Holroyd, C.B., 2016. Exploring individual differences in task switching: Persistence and other personality traits related to anterior cingulate cortex function. Prog. Brain Res. 229, 189–212.

Umemoto, A., Holroyd, C.B., 2017. Neural mechanisms of reward processing associated with depression-related personality traits. Clin. Neurophysiol. 128, 1184–1196. DOI:10.1016/j.clinph.2017.03.049

Van den Berg, I., Franken, I.H.A., Muris, P., 2010. A new scale for measuring reward responsiveness. Front. Psychol. 1, 1–7. DOI:10.3389/fpsyg.2010.00239

Vassena, E., Holroyd, C.B., Alexander, W.H., 2017. Computational models of anterior cingulate cortex: At the crossroads between prediction and effort. Front. Neurosci. 11, 1–9. DOI:10.3389/fnins.2017.00316

Verguts, T., Vassena, E., Silvetti, M., 2015. Adaptive effort investment in cognitive and physical tasks: a neurocomputational model. Front. Behav. Neurosci. 9. DOI:10.3389/fnbeh.2015.00057

Vogt, B.A., 2009. Cingulate Neurobiology and Disease. Oxford University Press, Oxford, UK.

Walsh, M.M., Anderson, J.R., 2012. Learning from experience: Event-related potential correlates of reward processing, neural adaptation, and behavioral choice. Neurosci. Biobehav. Rev. 36, 1870–1884. DOI:10.1016/j.neubiorev.2012.05.008

Wascher, E., Rasch, B., Sanger, J., Hoffmann, S., Schneider, D., Rinkenauer, G., Heuer, H.,Gutberlet, I., 2014. Frontal theta activity reflects distinct aspects of mental fatigue. Biol. Psychol. 96, 57–65. DOI:10.1016/j.biopsycho.2013.11.010

Westbrook, A., Braver, T.S., 2015. Cognitive effort: A neuroeconomic approach. Cogn. Affect. Behav. Neurosci. 15, 395–415. DOI:10.3758/s13415-015-0334-y

Wylie, G.R., Genova, H.M., DeLuca, J., Dobryakova, E., 2017. The relationship between outcome prediction and cognitive fatigue: A convergence of paradigms. Cogn. Affect. Behav. Neurosci. DOI:10.3758/s13415-017-0515-y

